# Fecundity Estimation of Atlantic mudskipper *Periophthalmus barbarus* in Ogbo-Okolo mangrove Forest of Santa Barbara River, Bayelsa State Niger Delta, Nigeria

**DOI:** 10.1101/2024.02.01.578404

**Authors:** Ayibatonyo Markson Nathaniel, Ilemi Jennifer Soberekon, Igoniama Esau Gamage, Akayinaboderi Augustus Eli, Morufu Olalekan Raimi

## Abstract

**Rationale:** Fecundity estimation and reproductive biology of Atlantic mudskippers *(Periophthalmus barbarus)* in the Niger Delta region of Nigeria needs to be studied.

**Objectives:** To estimate fecundity, gonadosomatic index, and condition factor of *P. barbarus* and describe its reproductive biology.

**Methods:** *P. barbarus* specimens were collected from Ogbo-Okolo mangrove forest in Bayelsa State, Nigeria. Length, weight, and gonad weight measurements were taken. Fecundity was estimated by the gravimetric method. Length-weight relationship, condition factor, and gonadosomatic index were calculated. Ovarian developmental stages were identified.

**Results:** Highest mean fecundity of 9612.7 eggs was observed in females of 10.1-12.0 cm standard length and 20.0-27.9 g weight. Length-weight relationship showed specimens were in good condition. Gonadosomatic index was higher in smaller individuals. Four ovarian developmental stages were identified.

**Conclusion:** *P. barbarus* exhibits high fecundity. Reproductive potential is greater in intermediate sized individuals compared to smaller or larger fish.

**Recommendations:** Sustainable management practices should be implemented to conserve *P. barbarus* stocks in the Niger Delta region. Further research into reproductive behavior and ecology is needed.

## Introduction

The Atlantic mudskipper *(Periophthalmus barbarus)* is a species of mudskipper native to the tropical Atlantic coasts of Africa, the Indian Ocean, and the western Pacific Ocean. They inhabit fresh, marine, and brackish waters, including tidal flats and mangrove forests, where they readily move across mud and sand out of the water. Mudskippers are members of the genus Periophthalmus, which includes oxudercine gobies characterized by dorsally positioned eyes and pectoral fins that aid in locomotion on land and in water. They skip, crawl, and climb using their pelvic and pectoral fins. As semi-aquatic animals, mudskippers transition between aquatic and terrestrial environments. They are carnivorous and utilize an ambushing strategy to capture prey using their hydrodynamic tongue to suction food into their mouths. Sexual maturity is reached at approximately 10.2 cm for females and 10.8 cm for males, and they can live for about five years. Mudskippers are used by people for food, bait, and medical purposes. They are primarily found in West African mangrove swamps and brackish water bodies along the coast (Ezenwari and Offiah, 2003; Hossain *et al*., 2011; Olalekan *et al*., 2019; Raimi *et al*., 2019; Raimi *et al*., 2022a; Saliu *et al*., 2023; Sylvester *et al*., 2023). The Atlantic mudskipper is found in several African countries including Angola, Democratic Republic of Congo, Cameroon, Nigeria and Ghana. Their distribution within these regions is influenced by the availability of food and shelter. Additionally, their hibernation capacity may impact their geographic distribution.

The scientific name *Periophthalmus barbarus* originated from the Greek words *‘peri*’ meaning ‘around’ and *‘ophthalmos’* meaning ‘eyes,’ referencing the eyes positioned dorsally that provide a large field of vision. In Greek, *‘barbarus’* means foreign, likely named for the mudskipper’s unusual characteristics compared to other gobies. The common name ‘mudskipper’ refers to their skipping movement across mudflats. They belong to the oxudercinae gobies, which inhabit both land and water. Mudskippers dig burrows for shelter and reproduction.

Initially, oxudercinae was described as a monotypic family, with all members classified under the single species Oxuderces dentatus. Oxudercinae are small to medium in body size, with elongated bodies covered in smooth scales. They can be identified by their dorsally positioned eyes and pointed, canine-like teeth. Their dorsal, pectoral, and pelvic fins contain spines, varying in number. Within oxudercinae, the Periophthalmus genus comprises 12 species distinguished by having teeth in a single row on the upper jaw and a maximum of 16 spines on the pectoral fins. All Periophthalmus live in mangroves or mudflats. *P. barbarus* is further identified by spots or white spots on the back and over 90 scales along the sides.

*P. barbarus* grows up to 16 cm long. Its scale-covered body is coated in mucus to retain moisture. It has over 90 scales along the sides and stores water in its gill chambers, allowing breathing out of water. The gill chambers lack a membrane cover and can open/close through surrounding muscles or differences in partial pressure (Michel *et al*., 2016). In addition to retaining moisture by storing water from the surface which helps them to breathe through its skin, otherwise known as cutaneous respiration (Kutschera and Elliot, 2013). They have pair of caudal fins that aid in aquatic locomotion and pelvic fins in terrestrial locomotion (Pace and Gibb, 2009). Their pelvic fins are adapted to terrestrial-living by acting as a sucker to attach on land, their eyes are adapted closely together and can move independently of about 360 degrees, their eyes are also positioned further up on the head, enabling the eyes to remain above the water surface while their body is submerged underwater (El-Sayed, 2006; Ansari *et al*., 2014). Cup-like structures that hold water are located beneath the eye which aids in lubricating the eyes when it is on land. They have chemosensory receptors that are located within the nose and on the skin surface (Etim, 2002; Chukwu and Deekae, 2011).

While, mudskippers have a mouth that can reorient. Their short digestive system comprises an esophagus, stomach, intestine and rectum (Wolczuk *et al*., 2018). The intestinal surface is folded, increasing the surface area for enhanced nutrient absorption. They have a unique olfactory organ including a 0.3 mm diameter canal near the upper lip that expands into a chamber-like sac. Genital papillae located on the abdomen differentiate females, which have less rounded papillae, from males (Kuciel, 2013). As semi-aquatic animals, mudskippers inhabit slightly salty waters such as rivers, estuaries and mudflats. They spend most daylight hours on land in tidal regions, emerging at low tide to feed and hiding in burrows at high tide. These burrows can extend 1.5 meters deep providing refuge from predators (Ansari *et al*., 2014). The burrows may contain air pockets for breathing despite low oxygen. Mudskippers tolerate high concentrations of industrial wastes like cyanide and ammonia in their environment (Emuebie, 2011; Siddik *et al*., 2016). When exposed to high ammonia, they can actively secrete it through their gills even in highly acidic conditions (pH=9.0) (Ansari *et al*., 2014). They survive varied environments including different water temperatures and salinities. Hot, humid climates optimally enhance cutaneous respiration and maintain their surface body temperature between 14-35°C.

Mudskippers build mud walls around their approximately 1-meter territory to maintain resources from predators. While hunting, they submerge leaving only their eyes out to sight and locate prey. They then launch onto land using predominantly their pectoral fins and catch prey in their mouth. When land predators threaten, they exhibit flight behavior by jumping into water or skipping away on mud (Ansari *et al*., 2014). Therefore, the objectives are to assess the length-weight relationship and condition factor of *P. barbarus*, and determine its gonadosomatic index.

## Materials and Methods

### Study Area

The Ogbo-okolo mangrove Forest of Santa Barbara River is located in Nembe local government Area of Bayelsa State, Nigeria at 4.5328°N, 6.4037°E (see figure 1 below). The area lies entirely below sea level with a maze of meandering creeks around the mangrove forest. Ogbo-okolo mangrove Forest of Santa Barbara River is significant in the provision of suitable breeding sites for diverse aquatic organism that abound in the area, good fishing ground for artisan fishers as well as petroleum exploration and production activities by Aiteo company.

**Figure 1.**
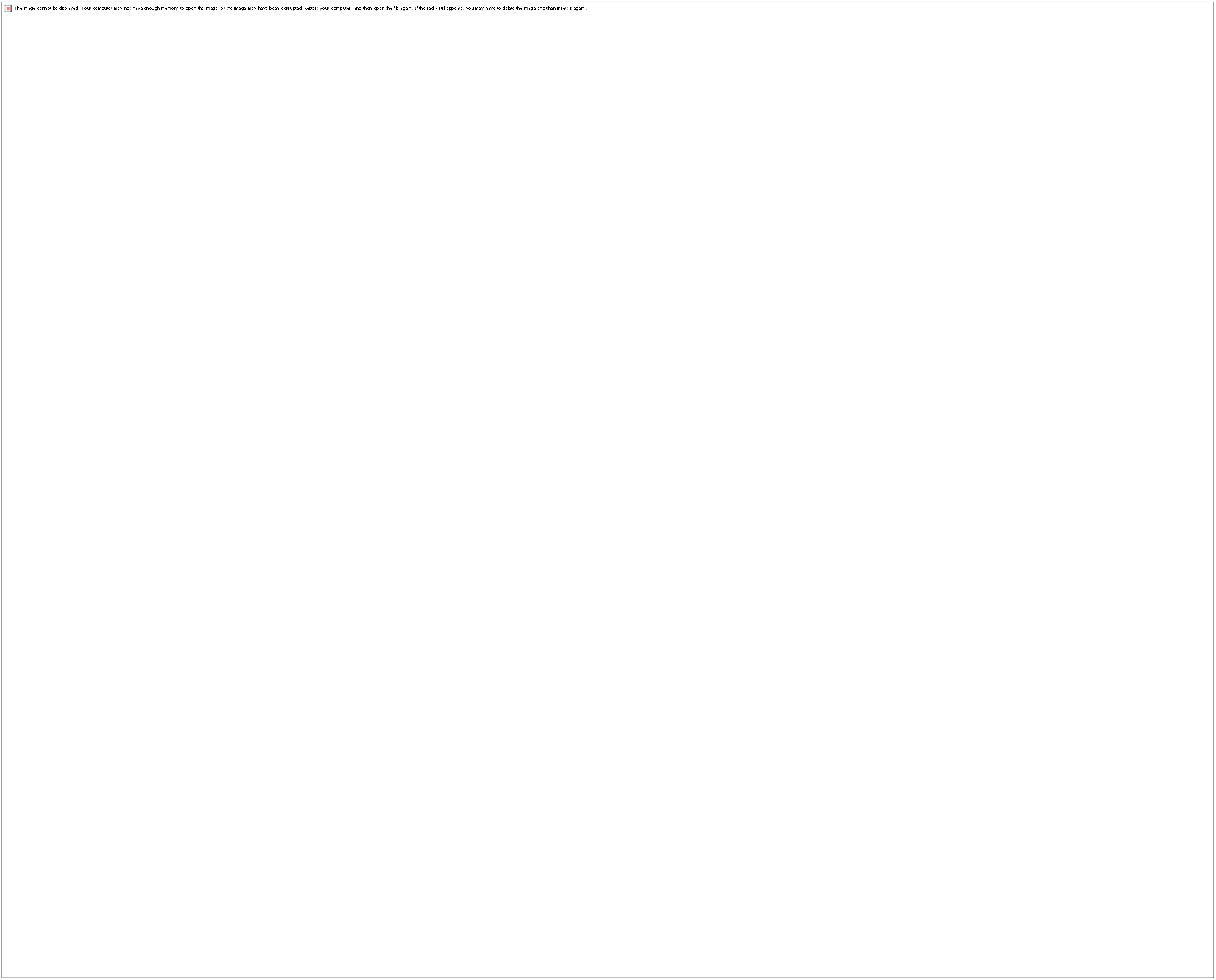
: Map of Nembe showing Santa Barbara River and the Study Area Ogbo-okolo mangrove Forest.

### Study site

The vegetation of the Ogbo-okolo mangrove consists of the red mangrove Rhizophora racemosa and white mangrove Avicenna Africana ranging in height from 5 to 15 meters. The main sea flows into smaller tributaries during high tide when the water is salty. At low tide, the stilt-like prop roots of mangroves are visible and the intertidal mudflat is exposed, serving as a feeding ground for mudskippers. The mudskippers dig small burrows between 3.5-6 cm in diameter around the prop-roots of Rhizophora trees. These areas serve as hideouts for *Periophthalmus* species. Traps will be set around these burrows to catch *Periophthalmus* specimens. Bacteria and microorganisms thriving in the mud produce sulfur gases, giving the mudflat a rotten egg odor. The study area has a tropical climate with average temperatures around 25°C and high humidity. Rainfall is heavy year-round, with peak precipitation from May to July averaging over 300 mm monthly as describe by studies from the Niger Delta region of Nigeria (Olalekan *et al*., 2022a, b; Raimi and Sawyerr, 2022; Raimi *et al*., 2022b, c, d; Glory *et al*., 2023; Raimi *et al*., 2023). The abundant rainfall contributes to the mangrove swamps and muddy conditions ideal for mudskippers.

### Sample Collection

Fish trapping method were used to collect 350 Specimen of Mudskippers *Periophthalmus barbarus* with standard length ranging from 5.5-11.9cm and total length ranging from 7.0-14.7cm respectively. The specimen was obtained in May 2021. The fishing gears used is hands made rubber container and basket traps woven with cane materials with a single conical in curved opened. Each basket trap was 40cm long and 30cm wide. These traps are nonselective and can catch both adults and Juveniles. The traps were set during the low tide using scattered and broken crabs likes *Callinetes sapidu* (marine swimming crab), *Cardiosoma armatum* (terrestrial crab), and *Sesarma huzardi* (hairy mangrove crab) was used as bait for the traps. As soon as catches were made, the specimens were removed and put in a bucket containing 5% formalin and a little water. These were later taken to the laboratory.

Standard length and total length of each sample was measured to the nearest 0.1 cm using a measuring board and the gutted weight of corresponding sample were measured to the nearest 0.1g. Specimens were dissected and the guts were carefully removed with the aid of a forceps after dissection. The gut and the liver were weighed together to the nearest 0.1g. The standard length, gutted weight and body weight of fishes were recorded into a data sheet for data analysis. In the analysis, the Total Length (TL) of the fish was measured from the tip of the anterior the (mouth part) to the caudal fin using an improvised measuring board made from wood. Fish weight was measured with an electronic scale, to the nearest 0.1g. The mean lengths and weights of the classes were used for data analysis. The relationship between the Standard Length (SL), TL, and total body weight (W)g of the fish is expressed by the equation below:

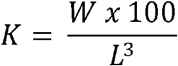

Where K = condition factor

W= weight of fish (g)

L = Total length (TL) of fish in (cm)

While, determining the Gonado somatic index of *Periophthalmus barbarus*. Biometric data of SL, TL to the nearest 0.01cm and body weight measurement (TW) and Gutted Weight (GW) to the nearest 0.01g were recorded. Further laboratory analysis was carried out by opening the stomach of the specimens to determine the opvaris of *Periophthalmus barbarus* to ascertain the gonad stages by use of naked eye examination of gonad development stage. Six biological stages of gonad development were identified as cited by Bagenal (1973). Moreover, four stages of gonad development were identified in this study and the classification of gonadal stages using Udo, (2002) methods. Each specimen ovaries and specimen gut were measured to the nearest 0.1g. Gonado somatic index (GSI) of the specimen was calculated according to El-Sayed *et al*., (2007) method:

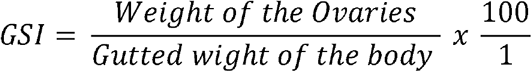

To determine the fecundity of Atlantic mudskipper *Periophthalmus barbarus* an unbiased sample of the specimen were obtained. Gonad stage IV and V were used in the fecundity estimation. The specimen with gonad stage IV and V were sorted out and the standard length measured to the nearest 0.01cm and gonad weight to the nearest 0.01g, the ovaries of each is removed and stored in specimen bottle containing 3% formalin and later place in plastic petri-dishes which were agitated at interval to ensure the eggs were separated from their connective tissue and spread after adding little water in the petri-dishes. Fecundity was estimated by taking the weight of the egg follow by the weight of subsample by Bagenal (1973) to the nearest 0.01g.

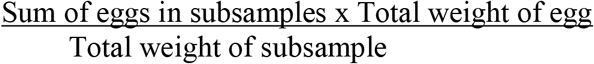

## Result

### The assessment of the length-weight relationship of Atlantic Mudskipper *P. barbarus* is shown below

The standard length ranging from 5.0-11.9cm SL, total length ranging 7.0-14.9g TL and the body weight between 3.4 - 30.6g BW

Condition factor:

The parameters of length-weight relationship are influenced by a series of condition factors including season, habitats, gonad maturity, sex, diet or available foods, stomach fullness, health of the individuals in their natural habitats as well as the treatment of specimens and preservation techniques.

The condition factor shows the good condition of the fish in terms of physical capacity for survival and reproduction. The condition factor was calculated to assess fish health in general, productivity and physiological conditions of fish populations. The results of the condition factor analyzed were standard length ranging from 5.0-7.0 and a total length ranging from 7.0 - 9.0 with weight ranging from 3.4-10.0 have condition factor of 3.0 and a standard length ranging from 8.1-9.0 and a total length ranging from 10.1 - 11.0 with weight ranging from 15.1-20.0 have condition factor of 2.8 and standard length ranging from 10.1-12.0 and a total length ranging from 12.6 - 14.9 with weight ranging from 25.1.30.9 have condition factor of 1.6, full detail shown in table 2 above which indicates that condition factor decrease with increase in size of the fish.

**Table 1:** Shows the length – weight relationship in *P. barbarus*.

| Standard Length (cm) |  | Total Length(cm) |  | Total Weight (G) |  |
| --- | --- | --- | --- | --- | --- |
| Range | Mean | Range | Mean | Range | Mean |
| 5.0-7.0 | 6.3 | 7.0-9.0 | 8.7 | 3.4-10.0 | 7.5 |
| 7.1-8.0 | 7.7 | 9.1-10 | 9.5 | 10.1-15.0 | 11.6 |
| 8.1-9.0 | 8.5 | 10.1-11.0 | 10.8 | 15.1-20.0 | 17.5 |
| 9.1-10.0 | 9.5 | 11.1-12.5 | 11.3 | 20.1-25.0 | 22.3 |
| 10.1-12.0 | 11.8 | 12.6-14.9 | 13.6 | 25.1-30.9 | 27.1 |

**Table 2:**
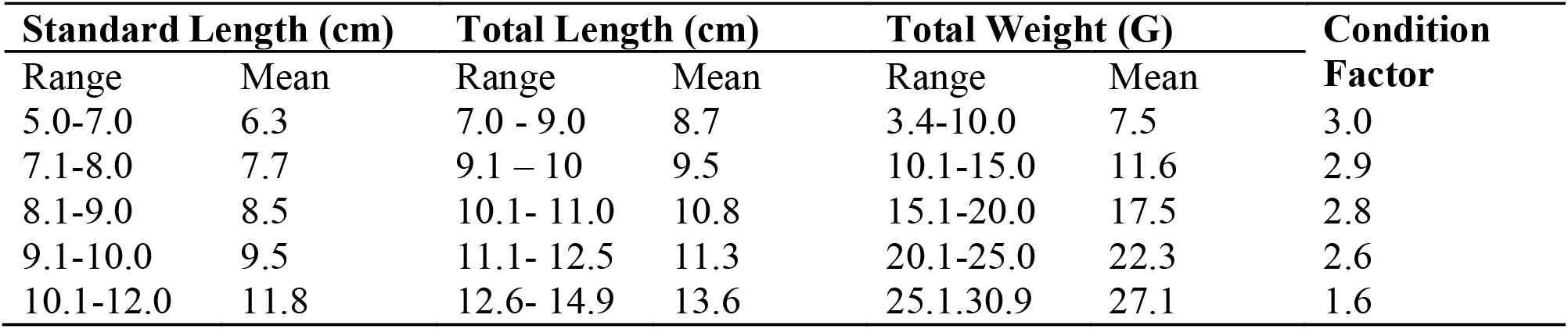
Relationship of length and weight to condition factor in *P. barbarus*.

In this study, four (4) Ovaries stages were observed in *P. barbarus*. These were Stages II - early development, stage III – late development, stage IV – matured, Stage V – Ripe or Gravid. Such table shows the macroscopic feature of the stages of ovaries development of *P. barbarus* and it revealed that medium size fish have the highest number of fish with ripe eggs (fecundity).

Gonadosomatic index (GSI) of *Periophthalmus barbarus* in table 4 revealed that gonado somatic index is lower in body weight ranging from 21.0 to 27.9g with 17.78 and higher in body weight ranging from 1.0-10.0g with 22.06. While mean of ovary weight 0.28 is lower in body weight ranging from 1.0 -10.0 and mean of ovary weight 0.45 is higher on the body weight ranging from 21.0-27.9. This indicates that GSI increase with body weight but decrease in mean of ovary weight.

**Table 3:** Relationship of length and body weight to stages of Gonad Maturation in *P. barbarus*.

| Number of Specimen Examined | Length (cm) |  | Total Body Weight (G) | Stages of Maturation |
| --- | --- | --- | --- | --- |
|  | Standard Length Range | Total Length Range |  |  |
| 4 | 7.0 - 8.0 | 8.0 -10.0 | 4.0 - 10.0 | **II** |
| 19 | 8.1 - 9.0 | 10.1 - 11.0 | 10.1 - 15.0 | **III** |
| 18 | 9.1 - 10.0 | 11.1 - 12.0 | 15.1 - 20.0 | **IV** |
| 5 | 10.0 – 12.0 | 12.1 – 14.0 | 20.1 - 27.5 | **V** |

**Table 4:** Gonadosomatic Index (GSI)

| Body weight Range (g) | Mean of ovary weight(g) | GSI |
| --- | --- | --- |
| **1.0 - 10.0** | 0.28 | 22.06 |
| **10.0 - 15.0** | 0.29 | 21.80 |
| **15.1 - 20.9** | 0.37 | 18.32 |
| **21.0 – 27.9** | 0.45 | 17.72 |

Table 5 showed that fish with standard length range from 9.1 - 10.0cm and weight ranging from 15.1 - 20.0g have the highest mean fecundity of 9612.7eggs but lower in standard length ranging 7.0 - 9.0cm with mean fecundity of 7428.8.

**Table 5:** Relationship of Fecundity in *P. barbarus* to length and body weight.

| Number of Specimen Examined | Length in (cm) |  | Total Body Weight (G) | Fecundity |  |
| --- | --- | --- | --- | --- | --- |
|  | Standard Length Range | Total Length Range |  | Mean | Range |
| 3 | 7.0 - 9.0 | 8.0 -10 | 4.0 - 15.0 | 7428.8 | 741.2- 9766.3 |
| 4 | 9.1 - 10.0 | 11.1 - 12.0 | 15.1 - 20.0 | 9612.7 | 5484-14570.5 |
| 3 | 10.0 – 12.0 | 12.1 – 14.0 | 20.1 - 27.5 | 8070.6 | 5048-12818 |

### Discussion

Length-weight relationship of Atlantic mudskipper, *Periophthalmus barbarus* were standard length ranging from 5.0-11.9cm SL, total length ranging 7.0-14.9g and the body weight between 3.4 - 30.6g body weight (BW). The condition factor values greater than one indicate the Periophthalmus barbarus specimens were in good physical condition. Condition factor decreased as fish size increased (Tables 1 and 2), aligning with the finding that condition factor is inversely related to length. The estimated allometric coefficient (b) from the length-weight relationships fell within the expected range of 2.0 to 4.0 reported for most fish (Froese and Pauly, 2006). The b value reflects that weight increases curvilinearly relative to length. Overall, the length-weight relationship parameters estimated in this study for *P. barbarus* were consistent with typical ranges reported for other fish species. This provides evidence that the sampled population had normal growth patterns and physiological condition. However, Siddik *et al*., (2016) reported isometric growth of *A. bato* from a southern coastal river of Bangladesh. But varies with the findings in Lagos lagoon, mudskipper, *Periophthalmus papilio* which was grouped into unsex, males and females with size ranging from 30.0-190 mm TL and weighing between 0.5 and 65.3g BW. There was size variation across the three groups through unsex, 32-158mm; males, 69-180mm and females, 60-190mm total length. Females attained higher growth and maturity at 69mm TL and males at 60mm TL in the study, males were significantly heavier in weight but shorter in body length than the females. Lawson (2011) also contradicted with 102mm and 105mm total length at maturity which he reported for females and males respectively. Udo, (2002) also reported isometric (b is equal to 3) length weight relationship for both sexes of this species. However Abdoli *et al*., (2009) estimated values is between 2.10 and 2.86 for both sexes for three species of mudskipper in their study in the coastal areas of the Persia Gulf in Iran. Similar trends were observed in the specie by Lawson, (2011) who reported values between 2.56 and 3.50 in the coastal areas of Selangor, Malaysia. A positive correlation values of r = 0.9385 (unsex), 0.9684 (males) and 0.9784 (females) showed there was a strong correlation between the total length and body weight measurements of the fish, meaning the fish increase with body weight as it grows in total length. This may be more genetical than being ecological. Strong correlation between fish body size and otolith weight of a related species *Periophthalmodon schlosseri* was also reported by (Sarimin *et al*., 2009). The parameters of length-weight relationship are influenced by a series of factor including season, habitat, gonad maturity, sex, diet, stomach fullness, health of the individuals in their natural habitats as well as the treatment of specimens and preservation techniques the condition factor of *P. barbarous* were found to be greater than one (1) which indicates that the specimen were relatively healthy. However, this study showed that condition factor decreases with increase in size of the fish. The naked eye examinations of the gonads of *P. barbarous*, forty-seven (47) specimen that have eggs, four stages of gonad development were identified in this study. These includes Stage II Early developing or virgin stage ll which ovary is very small firm and pale-white, stage III Late developing or maturing stage - ovary is small graining orange-yellow heavy network of vessel appear laterally on the surface of ovary wall, stage IV matured stage - ovary is yellow, large and selling the body cavity, ripe or gravid stage – ovary is yellow, large with contour walls turgid, distend body cavity, eggs are clearly distinct. Gonadosomatic index (GSI) is higher in body weight ranging 1.0 to 10.0g with 22.06 (GSI) and lower in body weight ranging from 21.0 to 27.9g with 17.78 (GSI) increase with increase in body weight but decrease in mean of ovary weight (Table 4). This showed that Atlantic mudskipper *Periophthamus barbarus* is highly productive between May and June in the study area this is contradicted with observation made by Atiqullah *et al*., (2013) on *Pomadasys stridens* which range from 0.801 to 9.124. Lawson, (2011) who reported that (GSI) values of the fish varied between 0.01 and 0.48% in males and 0.11-8.40% in females. Higher GSI values indicate a better wellbeing for the fish, this difference in values can be attributed to the combination of one or more factors including habitat, area, season, gonadal condition, sex, health, preservation methods and differences in the size and type of the specimens caught. fecundity in *P. barbarus* (Table 5) showed that fish with standard length range from 9.1 - 10.0cm and weight ranging from 15.1 - 20.0g have the highest mean fecundity of 9612.7eggs, but specimens range from 10.1-12.0 standard length and weight ranging from 20.0-27.9g are more fecund than those smaller or bigger than the bracket which have fewer eggs. This means fecundity estimated for the fish with this size range 9.1 to 10.0cm TL and 15.1 to 20.8 g W, are more fecund. This showed higher fecundity contributed to body weight. The fecundity of the species increased with fish length and body weights. This is in line with an observation made by Mfon and King (2001) who suggest the egg production capacity of the gobiid mudskipper.

### Conclusion

The study on Atlantic mudskippers, *P. barbarus*, in the Ogbo-okolo mangrove forest demonstrated that reproductive output is associated with body size. Specimens with a standard length range of 9.1-10.0 cm and weight 15.1-20.0 g exhibited the highest mean fecundity at 9,612.7 eggs. Fish within the size range of 10.1-12.0 cm standard length and 20.0-27.9 g weight showed greater fecundity compared to smaller individuals, indicating that higher fecundity accompanies larger body weight. Overall, the fecundity of *P. barbarus* increased with fish length and body weight, which aligns with trends observed for many fish species. Larger females tend to produce more eggs due to having greater energy reserves for gamete production. The higher fecundity of intermediate sized *P. barbarus* may reflect an optimal balance of energy allocation for growth versus reproduction. Smaller individuals may invest more in somatic growth, while larger fish experience diminished returns on egg production. This study provides baseline data on the reproductive biology of *P. barbarus*, an important component of mangrove ecosystems in the Niger Delta region. The findings can inform sustainable management efforts for this species. Further research over broader geographic and temporal scales would clarify drivers of variability in fecundity and other reproductive parameters. Investigating the relationship between female body condition, environmental factors, and reproductive output could provide additional insights into the ecology of *P. barbarus*.

### Policy Implication

Atlantic mudskippers (*P. barbarus*) are economically and ecologically important components of Niger Delta mangrove ecosystems. This study found the fecundity and reproductive potential of *P. barbarus* populations in Bayelsa state is connected to adult body size, with implications for sustainable management. Overfishing and habitat loss threaten the sustainability of this fishery. To prevent declines in fish stocks, reproductive output, and future harvests, policy makers should implement science-based regulations on P. barbarus fishing. Required actions include:

- Set minimum size limits to allow females to attain optimal size for reproduction before capture
- Restrict harmful gears like dynamite fishing which indiscriminately kill juveniles and breeding adults
- Designate protected mangrove areas as essential fish breeding habitat
- Support mangrove restoration to mitigate habitat losses
- Provide education programs and incentives for sustainable fishing practices
- Improve monitoring and enforcement to ensure compliance

Urgent efforts are needed to balance economic demands for P. barbarus with maintenance of its ecological role. Sustainable management informed by research will provide long-term social, economic, and environmental benefits.

### Recommendations

Based on the findings from this research work, the following recommendations were made:

- Additional research on *P. barbarus* should be conducted across its full geographic range in Bayelsa state using standardized methods to allow for broader comparisons of reproductive parameters and population dynamics. Studies across multiple seasons would provide greater insight into temporal variability.
- Sustainable management plans for *P. barbarus* should be developed and implemented by fisheries regulators in collaboration with local communities. These may include size limits, gear restrictions, spatial protections, and harvest quotas to prevent overexploitation.
- Educational programs should be created to inform fishermen about sustainable fishing practices for *P. barbarus* and the importance of this species to the mangrove ecosystem. This knowledge can encourage voluntary adoption of responsible harvesting techniques.
- Mangrove habitat restoration efforts in areas degraded by pollution or development pressures are recommended to preserve critical nursery and breeding grounds for *P. barbarus*.

## Competing Interests

All authors have declared no competing interests exist.

## References

Abdohi, L. E., Kamrani, A. Abdoli and Kiabi B. (2009) Length weight relationships for three species of mudskippers (Gobndae: Oxudercinane) in the coastal areas of the Persian Gulf, Iran. J Applied Ichthyol. 52:236–237.

Ansari, A., Trivedi S, Saggu S, Rehman, H. (2014). Mudskipper: A biological indicator for environmental monitoring and assessment of coastal water. Journal of Entomology and zoology studies, 5481–6035.

Atiqullah, M., Hossain, M. M., Kamal, M. S., AllJHarthi, M. A., Khan, M. J., Hossaen, A., & Hussain, I. (2013). Crystallization kinetics of PElJblJisotactic PMMA diblock copolymer synthesized using SiMe 2 (Ind) 2 ZrMe 2 and MAO cocatalyst. AIChE Journal, 59(1), 200–214.

Bagenal, T.B (1973) Fish fecundity and its relations with stock and recruitment. Rep. P.V. Renn. Cons Int. Explore Mer., 164:186–198.

Chukwu, K. and Deekae, S., (2011). Length-weight relationship, condition factor and size composition of Periophthalmus barbarus (Linneaus 1766) in New Calabar River, Nigeria. Agriculture and Biology Journal of North America, 2(7), 1069–1071.

El-Sayed KH, Moharrams G. (2007) Reproductive biology of Tilapia zilli (Gerv, 1848) from Abu Qir Bay, Egypt. Egyptian Journal of Aquatic Research; 33(1):379–394.

El-Sayed AFM (2006). Tilapia culture. CABI Publishing, Wallingford OX 108 DE. UK. 273.

Emuebie Okonj Raphael (2011) Physio Chemical Properties of Mudskipper (Periophthalmus barbarus (pallas) liver Rhodanese” Australian Journal of Basic and Applied Sciences 5(8), 507–514.

Etim, Lawrence (2002) Breeding, Growth, Mortality and Yield of the Mudskippers Periophthalmus barbarus (Teleostei: Gobiidae) in Imo River estuary, Nigeria 227–238.

Ezenwaji H.M.G., & Offiah F.N., (2003) Reproductive biology of Periophthalmus barbarus in Calabar River, Nigeria, 1158–1161.

Froese R, Pauly D., (2006). Fish Base. World Wide Web electronic publication, http://www.fishbase.org.

Glory Richard, Sylvester Chibueze Izah, Morufu Olalekan Raimi and AustinAsomeji Iyingiala (2023) Public and environmental health implications of artisanal petroleum refining and risk reduction strategies in the Niger Delta region of Nigeria. Journal of Biological Research & Biotechnology. Vol. 21 No.1; pp. 1836–1851. ISSN (print):1596-7409; eISSN (online):2705-3822. 10.4314/br.v20i3.6. http://www.bioresearch.com.ng. Publisher: Faculty of Biological Sciences, University of Nigeria, Nsukka, Nigeria.

Hossain M Y, Rahman M M, Fulanda B, Jewel MAS, Ahamed F, Ohtomi J. (2011). Length-weight and length-length relationships of five threatened fish species from the Jamuna Brahmaputra River Tributary River, northern Bangladesh. Journal of Applied Ichthyology. 28, 275–277.

Kuciel, Michel (2013) The mechanism of olfactory organ ventilation in Periophthalmus barbarus (Go biidaeoxudercnae. zoo morphology. 132 (1), 81–85.

Kutschera U & Elliot M., (2013). Do mudskippers and lungfishes elucideta the early evoluto of four-limbed vertebrates? Evolution and outreach. 6(1), 8.

Khaironizam, M.Z. and Norma Rashid, Y., (2002). Length-weight relationship of mudskippers (Gobiidae: Oxudercinae) in the coastal areas of Selangor, Malaysia. 25(3-4), 20–22.

Lawson E.O, (2011) testicular maturation and reproductive cycle in mudskipper, Periophthalmus papilio from Lagos lagoon, Nigeria. Journal of American Science 7(1), 48–59.

Lawson E.O (2011). Length-weight Relationships and Fecundity Estimates in Mudskipper, Periophthalmus papilio (Bloch and Schneider 1801) Caught from the Mangrove Swamps of Lagos Lagoon, Nigeria. Journal of Fisheries and Aquatic Science, 6, 264–271.

Michel, K. B, Aerts, P. and Van Wassenbergh, S. (2016) Environment-dependent Prey Capture in the Atlantic Mudskipper (Periophthalmus barbarus). Biol. Open 5, 1735–1742.

Mfon Timothy Udo & King R.P (2001) Fecundity of the mudskipper Periophthalmus barbarus (Gobiidae) in Imo River, Nigeria. Archive of Fishery and Marine Research 49(2),117–124.

Olalekan MR, Albert O, Iyingiala AA, Sanchez DN, Telu M (2022a) An environmental/scientific report into the crude oil spillage incidence in Tein community, Biseni, Bayelsa state Nigeria. J Environ Chem Toxicol. 2022;6(4):01–01., doi:10.37532/pulject.2022.6(4);01-06.

Olalekan MR, Olawale HS, Clinton IE and Opasola AO (2022b) Quality Water, Not Everywhere: Assessing the Hydrogeochemistry of Water Quality across Ebocha-Obrikom Oil and Gas Flaring Area in the Core Niger Delta Region of Nigeria. Pollution, 8(3): 751–778.

Olalekan RM, Omidiji AO, Williams EA, Christianah MB, Modupe O (2019). The roles of all tiers of government and development partners in environmental conservation of natural resource: a case study in Nigeria. MOJ Ecology & Environmental Sciences 2019;4(3):114lJ121. DOI: 10.15406/mojes.2019.04.00142.

Pace, C.M. Gibb, A.C. (2009) Mudskipper pectoral fin Kinematics in aquatic and terrestrial environments. Journal of Experimental Biology 212(14) 2279–2286.

Raimi, M.O., Oyeyemi, A.S., Mcfubara, K.G., Richard, G.T., Austin-Asomeji, I., Omidiji, A.O. (2023). Geochemical Background and Correlation Study of Ground Water Quality in Ebocha-Obrikom of Rivers State, Nigeria. Trends Appl. Sci. Res, 18(1), 149–168. 10.17311/tasr.2023.149.168.

Raimi MO, Abiola OS, Atoyebi B, Okon GO, Popoola AT, Amuda KA, Olakunle L, Austin AI & Mercy T. (2022a). The Challenges and Conservation Strategies of Biodiversity: The Role of Government and Non-Governmental Organization for Action and Results on the Ground. In: Chibueze Izah, S. (eds) Biodiversity in Africa: Potentials, Threats, and Conservation. Sustainable Development and Biodiversity, vol 29. Springer, Singapore. 10.1007/978-981-19-3326-4_18.

Raimi, MO., Sawyerr, HO., Ezekwe, IC., & Gabriel, S. (2022b). Toxicants in Water: Hydrochemical Appraisal of Toxic Metals Concentration and Seasonal Variation in Drinking Water Quality in Oil and Gas Field Area of Rivers State, Nigeria. In P. H. Saleh, & P. A. I. Hassan (Eds.), Heavy Metals - New Insights [Working Title]. IntechOpen. 10.5772/intechopen.102656. ISBN 978-1-80355-526-3.

Raimi, O., Ezekwe, C., Bowale, A. and Samson, T. (2022c) Hydrogeochemical and Multivariate Statistical Techniques to Trace the Sources of Ground Water Contaminants and Affecting Factors of Groundwater Pollution in an Oil and Gas Producing Wetland in Rivers State, Nigeria. Open Journal of Yangtze Oil and Gas, 7, 166–202. doi: 10.4236/ojogas.2022.73010.

Raimi OM, Sawyerr OH, Ezekwe CI, Gabriel S (2022d) Many oil wells, one evil: comprehensive assessment of toxic metals concentration, seasonal variation and human health risk in drinking water quality in areas surrounding crude oil exploration facilities in rivers state, Nigeria. International Journal of Hydrology. 2022;6(1):23lJ42. DOI: 10.15406/ijh.2022.06.00299.

Raimi, M. and Sawyerr, H. (2022) Preliminary Study of Groundwater Quality Using Hierarchical Classification Approaches for Contaminated Sites in Indigenous Communities Associated with Crude Oil Exploration Facilities in Rivers State, Nigeria. Open Journal of Yangtze Oil and Gas, 7, 124–148. doi: 10.4236/ojogas.2022.72008.

Raimi M O, Suleiman R M, Odipe O E, Salami J T, Oshatunberu M, et al (2019). Women Role in Environmental Conservation and Development in Nigeria. Ecology & Conservation Science; 1(2): DOI: 10.19080/ECOA.2019.01.555558. Volume 1 Issue 2 - July 2019. https://juniperpublishers.com/ecoa/pdf/ECOA.MS.ID.555558.pdf

Saliu, A.O., Komolafe, O.O., Bamidele, C.O., Raimi, M.O. (2023). The Value of Biodiversity to Sustainable Development in Africa. In: Izah, S.C., Ogwu, M.C. (eds) Sustainable Utilization and Conservation of Africa’s Biological Resources and Environment. Sustainable Development and Biodiversity, vol 888. Springer, Singapore. 10.1007/978-981-19-6974-4_10.

Sarimin, A. S., Ghaffar, M. A., Mohammed, C. A. (2009). Variation of Ca, Sr, Ba and Mg in the Otolith of Mudskipper in West Coast of Peninuslar Malaysia. Pak. J. Biol. Sci., 2009; 12:231–238.

Siddik MAB, Chaklader MR, Hanif MA, Islam MA, Fotedar R. (2016). Length-weight relationships of four fish species from a coastal artisanal fishery, southern Bangladesh. Journal of Applied Ichthyology. 1–3.

Siddik M, Hanif M, Chaklader M, Nahar A, Mahmud S, et al. (2016) Fishery biology of gangetic whiting Sillaginopsis panijus (Hamilton, 1822) endemic to Ganges delta, Bangladesh. Egyptian Journal of Aquatic Research 41: 307–313.

Sylvester CI, Odangowei IO, Matthew CO, Saoban SS, Zaharadeen MY, Muhammad A, Morufu OR, and Austin-Asomeji I (2023) Historical Perspectives and Overview of the Value of Herbal Medicine. In: Izah S. C, et al. (eds.), Herbal Medicine Phytochemistry, Reference Series in Phytochemistry, 10.1007/978-3-031-21973-3_1-1.

Udo M. T., (2002). Tropic attributes of the mudskipper, Periophthalmus barbarous (Gobiidae: Oxudercinae) in the mangrove swamps of Imo Riverestuary, Nigeria, African journal online 5(2), 508–517.

Wolczuk Katarzyna, Ostrowski Maciel, Ostrowska Angnieszka, Napiorkowska Teresa (2018) “Structure of the alimentary tract in the Atlantic Mudskipper Periophthalmus barbarous (Gobiidae: Oxudercinae): anatomical, historical and ultrastructural studies”. Zoology. 128: 38–45.

